# High-frequency quantitative ultrasound to assess the acoustic properties of engineered tissues in vitro

**DOI:** 10.1101/2022.08.03.502702

**Authors:** Joseph A. Sebastian, Eric M. Strohm, Emmanuel Chérin, Bahram Mirani, Christine Démoré, Michael C. Kolios, Craig A. Simmons

## Abstract

Acoustic properties of biomaterials and engineered tissues reflect their structure and cellularity. High-frequency ultrasound (US) can non-invasively characterize and monitor these properties with sub-millimetre resolution. We present an approach to estimate the acoustic properties of cell-laden hydrogels that accounts for frequency-dependent effects of attenuation in coupling media, hydrogel thickness, and interfacial transmission/reflection coefficients of US waves, all of which can bias attenuation estimates. Cell-seeded fibrin hydrogel disks were raster-scanned using a 40 MHz US transducer. Thickness, speed of sound, acoustic impedance, and acoustic attenuation coefficients were determined from the difference in the time-of-flight and ratios of the magnitudes of US signals, interfacial transmission/reflection coefficients, and acoustic properties of the coupling media. With this approach, hydrogel thickness was accurately measured by US, with excellent agreement to confocal microscopy (r^2^ = 0.97). Accurate thickness measurement enabled acoustic property measurements that were independent of hydrogel thickness, despite up to 60% reduction in thickness due to cell-mediated contraction. Notably, acoustic attenuation coefficients increased with increasing cell concentration (p<0.001), reflecting hydrogel cellularity independent of contracted hydrogel thickness. This approach enables accurate measurement of the intrinsic acoustic properties of biomaterials and engineered tissues to provide new insights into their structure and cellularity.

## 1 Introduction

Ultrasound (US) is an emerging measurement technique in biomaterials and tissue engineering (TE) applications due to its non-invasive, non-destructive, and real-time monitoring capabilities [1-5]. Beyond its well-known imaging capabilities, US can also be used to measure acoustic properties related to structural, functional, and intrinsic material properties of interest for biomaterials and engineered tissue applications. For example, US measurements of the speed of sound, acoustic impedance, and acoustic attenuation in a biomaterial correlate with the material’s microstructure [6], elastic properties [7], and cellularity [8, 9], respectively.

High frequency US (>10 MHz) is particularly attractive for in vitro TE applications, as it offers sub-millimetre spatial resolution to enable tissue- and cell-scale measurements (at tens and hundreds of MHz, respectively). However, a particular challenge with the application of US in these applications at high frequencies is that the bandwidth of the US transducers is large, the attenuation high, and the relationship between attenuation and frequency can become non-linear. Thus, an accurate estimation of the acoustic attenuation of biological media at high frequencies is critical to determining other acoustic properties (e.g., backscatter coefficient) and inferring the material properties of engineered tissues and cells.

In previous studies, attenuation in biological samples was estimated using a substitution method, whereby the US signals propagating between two transducers [7, 8, 10, 11], or reflected from a reference substrate (e.g., quartz) [12-14], were acquired in the presence and absence of the sample in the US path, and their amplitudes compared in either the time- and/or frequency-domain. However, as implemented in these studies, the method fails to fully compensate for the attenuation in the coupling medium, for the transmission of US waves at the coupling medium-biomaterial interface and, when required, for the reflection at the biomaterial-substrate interface. If not compensated for, transmission and/or reflection coefficients at interfaces introduce a bias in the estimate of the sample attenuation coefficient. This bias increases with the difference in acoustic impedance between the sample and coupling medium and is inversely proportional to the sample thickness. This bias becomes significant for attenuation measurements in thin samples and/or for significant differences in impedance between media. Furthermore, the sample thickness is often assumed based on sample preparation [8,9,11], which fails to account for dynamic reorganization of biomaterials by embedded cells that can alter thickness [15] and consequently affect the attenuation coefficient estimate. As an alternative to substitution methods, Ruland et al. [16, 17] recently introduced a reference phantom method (RPM) for quantitative US of cell-laden hydrogels and bioscaffolds. In the RPM, a sample’s acoustic properties are determined by comparison to a reference phantom of known attenuation and sound speed. However, this approach is limited to samples with similar acoustic properties to the RPM and also does not account for cell-mediated effects on the biomaterial shape and size and their effect on acoustic properties, and thus would be challenging to implement to track dynamic changes. Thus, compensation for transmission/reflection at interfaces and measurement of the biomaterial sample thickness is critical for accurately determining its acoustic attenuation coefficient and microstructural assessment.

Here, we present an US method of estimation of the thickness and acoustic properties of engineered tissues. We benchmarked this technique against published acoustic property values of polymethlypentene (TPX). This method accounts for the frequency- and thickness-dependent effects of attenuation in the coupling medium and reflection/transmission of US waves at interfaces. We demonstrate thickness-independent measurement of the speed of sound, acoustic impedance, and acoustic attenuation coefficient of fibrin hydrogels as a model biomaterial, at four different cell densities (cell-free control, 1×10^5^, 1×10^6^, or 1×10^7^ cell/ml) and three different initial thicknesses (1.25 mm, 1.00 mm, 0.75 mm). From the estimated acoustic properties, we derive the density and elastic modulus of the cell-laden hydrogels.

## 2 Methods

### 2.1 Acoustic characterization

Acoustic properties of samples were estimated at high frequency using a substitution method adapted from Briggs et al. [18] and depicted in Fig. 1A. US echoes reflected from the substrate surface (polystyrene in our case) with and without the sample inserted in the propagation path and reflected from the surface of the sample (*s*_2_(*t*), *s*_1_(*t*) and *s*_3_(*t*), respectively) are collected. Their Fourier transforms *S*_*i*_(*f*) (*i* = 1, … 3) are calculated to evaluate, in the frequency domain (Supplementary Figure 1), the coefficients of reflection (*R*_*w*−*s*_) and transmission (*T*_*w*−*s*_) at the water-sample interface, the coefficient of reflection (*R*_*w*−*p*_) at the water-substrate interface, and the coefficient of reflection (*R*_*s*−*p*_) at the sample-substrate interface. These coefficients are compensated for in equation (1), to get an unbiased estimation of the frequency-dependent attenuation in the sample *α*_*s*_ (Supplementary Figure 1):

**Figure 1:**
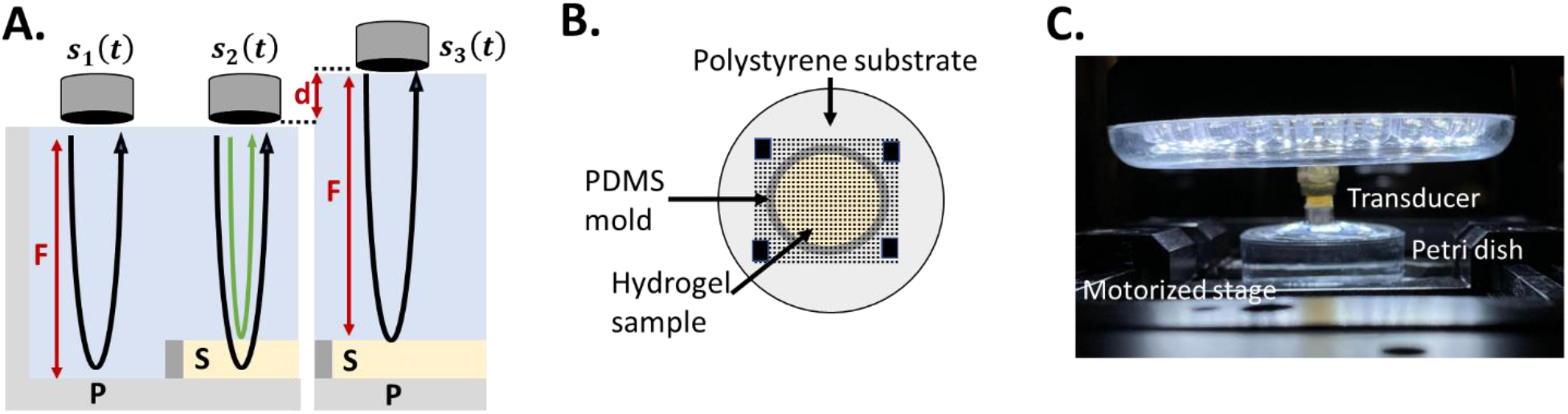
(A) Schematic representing the experimental system used to estimate the sample acoustic properties where S is the sample, P is the polystyrene substrate, F is the focal distance of the 40 MHz transducer, and d is sample thickness. *s*_1_(*t*) is the signal collected from the substrate surface (polystyrene in our case), *s*_2_(*t*) is the signal collected with the sample inserted in the propagation path, and *s*_3_(*t*) is the signal reflected from the surface of the sample. *s*_1_(*t*) and *s*_2_(*t*) are measured together (see description in B.) and *s*_3_(*t*) is measured separately. The thickness of the sample *d* is estimated from the difference in time of flight between the echo from the uncovered substrate *s*_1_(*t*) and the echo from the top of the phantom (green arrow) while focusing on the substrate. (B) Top view of a hydrogel sample cast in an 8 mm diameter polydimethylsiloxane mold positioned in the polystyrene Petri dish. The overlaid dotted lines represent the 15 mm x 15 mm raster scan with 100 µm step size in the x- and y-directions for the ultrasound measurements. The black squares represent the region where *s*_1_(*t*) was acquired. (C) Picture of the imaging system with the front face of the ultrasound transducer immersed into the polystyrene Petri dish filled with water.

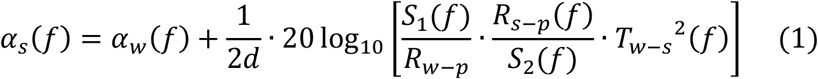

where:

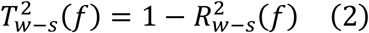

with:

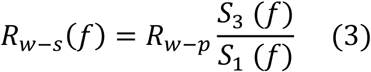

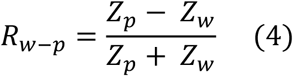

In equation (4), *Z*_*w*_ and *Z*_*p*_ are the acoustic impedances of water and the polystyrene substrate, respectively, and known. The reflection coefficient at the sample-substrate interface *R*_*s*−*p*_ in equation (1) is calculated from *Z*_*p*_ and the acoustic impedance of the sample *Z*_*s*_:

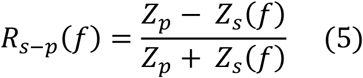

with:

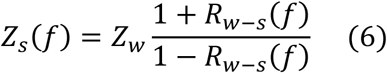

The thickness of the sample *d* is estimated from the difference in time of flight *Δt* between the echo from the uncovered substrate *s*_1_(*t*) and the echo from the top of the phantom (Fig. 1A, green arrow) while focusing on the substrate, and the speed of sound in water *c*_*w*_:

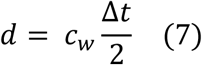

Time of flight was estimated as the times at which the peaks of the envelope of these echoes (*s*_1_(*t*) and *s*_3_(*t*)) were detected relative to US pulse transmission. Finally, since the logarithmic term in equation (1) corresponds, once normalized to the sample thickness, to the difference in attenuation between sample and water, water attenuation *α*_*w*_ is added to calculate the attenuation in the sample *α*_*s*_.

The speed of sound in the sample can be calculated from the thickness of the sample estimated in (7), and the difference in time of flight *Δt*′ between the echo from the surface of the sample (Fig. 1A, green arrow) and the echo from the underlying substrate *s*_2_(*t*):

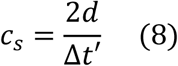

Finally, the sample density *ρ*_*s*_ and elastic modulus *ε*_*s*_ can be estimated from the sample acoustic impedance and speed of sound, and speed of sound and density, respectively:

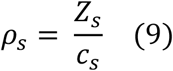

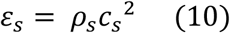

### 2.2 High-frequency ultrasound system and signal acquisition

A custom-built US system was developed to acquire ultrasound signals from the sample and substrate. This system comprises a single-element spherically focused 40-MHz transducer (focal distance F=8.56 mm, f-number 3) providing a 36 µm axial resolution, 110 µm lateral resolution (in the focal zone), and 4 mm depth-of-field (all measured at -6 dB) using a 40-µm aperture needle hydrophone (NH0040, Precision Acoustics Ltd, Dorchester, U.K.). The transducer was affixed to a microscope (IX71, Olympus America Inc., Melville, NY) equipped with a 3D-motion stage (Prior Scientific, USA; 2 motorized lateral axes X and Y, 1 manual vertical axis Z) on which the sample was positioned in a polystyrene Petri dish (Greiner Bio-One, North Carolina, USA), covered with water. The sample was moved relative to the transducer in a raster scan manner with a 100 μm step size in both lateral directions and such that a 15 mm x 15 mm region of interest (ROI). The ROI incorporated the sample, underlying substrate, and some uncovered substrate (Fig. 1B). US signals *s*_1_(*t*), *s*_2_(*t*) and *s*_3_(*t*) (Fig. 1A) were acquired at each lateral position (*x, y*), in two separate raster scans. For these acquisitions, monocycle pulses generated by a pulse generator (AVB2-C-OCIC, Avtech Electrosystems Ltd, Nepean, ON, Canada) were transmitted through a radiofrequency (RF) switch (Mini-Circuits, Brooklyn, NY, USA) to the transducer. Reflected ultrasound signals were collected by the same transducer, amplified by a 30 dB amplifier (Model # AU-2A-0150, Narda-MITEQ, Hauppauge, NY, USA) then digitized at 625 MS/s by 14-bit resolution A/D board (Teledyne SP Devices, Linköping, Sweden). At each location, signals were averaged 45 times to increase the signal-to-noise ratio and saved to a computer for offline processing and analysis using MATLAB (MathWorks, Natick, MA, USA). With a pulse repetition frequency of 10 kHz, the time to acquire data from a single raster scan was 8 minutes. The acquisition system hardware was computer-controlled using a trigger card (SpinCore Technologies, Gainesville, FL, USA). The entire system was enclosed in a temperature-controlled incubator (In Vivo Scientific LLC, Salem, SC, USA).

US data acquisition was performed with a thin polymethyl-pentene film (DX-845 TPX, C. S. Hyde Company, Lake Villa, IL, USA) and with hydrogel samples for validation of the acoustic characterization method (see below). Finally, cell-laden hydrogels were scanned and characterized. For TPX, the incubator temperature was set to 22°C, whereas for the hydrogel samples it was set to 37°C.

### 2.3 Method validation

To validate our method, the acoustic properties of a thin DX-845 grade TPX film were estimated and compared to those reported in [19] and [20]. The film was taped to a polystyrene Petri dish and covered with water at 22°C to match the temperature conditions of [19]. RF signals were collected as described in 2.2, and all material properties were calculated using equations (1) to (10), with *c*_*w*_ = 1488 m/s [20], *ρ*_*w*_ = 997.77 kg/m^3^ [21], *c*_*p*_ = 2410 m/s [22], *ρ*_*p*_ = 1050 kg/m^3^ (general purpose polystyrene) and resulting acoustic impedances *Z*_*w*_ = 1.485 MRayl and *Z*_*p*_ = 2.531 MRayl. For the attenuation in water in equation (1), which can be expressed as *α*_*w*_(*f*) = *af*^2^, *a* was set to 2.22 × 10^−4^ dB/mm/MHz^2^. For comparison, the film thickness was measured using a spherical digital tube micrometer (Mitutoyo Canada Inc., Mississauga, CA) before the acoustic measurement.

Since sample thickness plays a critical role in evaluating the acoustic properties, the acoustic measurement of the thickness of fibrin hydrogel samples was compared to laser confocal microscopy measurement. To fabricate the fibrin hydrogel, equal volumes of fibrinogen (Sigma-Aldrich, CAT# F8630) solution in Dulbecco’s phosphate-buffered saline (Gibco™ DPBS -/-, ThermoFisher Scientific, CAT# 14190144) and thrombin (Sigma-Aldrich, CAT# T6634) solution in complete cell culture medium were mixed to reach final concentrations of 5 mg/ml and 3 unit/ml for fibrinogen and thrombin, respectively. The mixture was cast in polydimethylsiloxane (PDMS) disk-shaped molds with a diameter of 8 mm (Fig. 1B) and initial thicknesses of 0.75 mm, 1.00 mm, and 1.25 mm. The mixture was incubated at 37 °C and 100% humidity for 90 min for crosslinking, after which DPBS +/+ (Corning, CAT# 21-030-CV) was added, and US measurements were conducted. Then, fibrin gels were stained by exposure to 50 µl of tetramethylrhodamine isothiocyanate (TRITC)–Dextran (Sigma-Aldrich – CAT# T1037) solution in DPBS +/+ at a concentration of 5 mg/ml and incubation at 37 °C for 1 hr. The hydrogels were washed using DPBS +/+. Their thickness was measured using a laser confocal microscope (Olympus FV3000) with emission at 640 nm by focusing on the lowest and highest surfaces where a meaningful signal was detected at a single location. The distance between these two surfaces was taken as the hydrogel sample thickness and compared with US measurements (eq. 7), for which the speed of sound in DPBS at 37°C was assumed to be that of pure water at the same temperature (*c*_*w*_ = 1524 m/s [21]). All the MATLAB processing codes, A-line data, statistical analyses, and selected ROIs are publicly available <u>here</u> for other researchers to reproduce the results of this study.

### 2.4 Cell-laden hydrogel acoustic characterization

Cell contraction can change the thickness and size of a cell-laden gel [13]. Although high-frequency US studies have examined cell-laden gels, cell contraction in hydrogels and engineered tissues have not been taken into account [7-11, 16, 17], despite the unavoidable effects on the physical and acoustic properties of these media. To address this issue, we seeded fibrin hydrogels, with initial thicknesses of 0.75, 1, and 1.25 mm, with cells at densities of 1×10^5^, 1×10^6^ and 1×10^7^ cells/ml.

Cell-laden hydrogels were prepared using human umbilical perivascular cells (hUCPVC) (Tissue Regeneration Therapeutics Toronto, Ontario, Canada). Cells at passage three were cultured in T175 flasks (CAT# 431080, Corning Life Sciences®, Tewksbury, Massachusetts, United States) in Minimum Essential Medium (MEM) Alpha (Gibco™ MEM α, CAT# 12561056, ThermoFisher Scientific, Mississauga, Ontario, Canada), at 37°C in an atmosphere of 5% CO2 and 100% humidity. MEM was supplemented with 20% v/v fetal bovine serum (Gibco™ FBS, CAT# 12483-020, ThermoFisher Scientific, Mississauga, Ontario, Canada), 0.24% v/v L-glutamine 200 mM (CAT# G7513, Sigma Aldrich, Oakville, Ontario, Canada), and 0.24% v/v penicillin-streptomycin 10,000 unit/ml (Gibco™, CAT# 15140122, ThermoFisher Scientific, Mississauga, Ontario, Canada). The culture medium was refreshed every three days. Then, hUCPVCs at passage four were suspended in the thrombin solution used in the hydrogel preparation described in section 2.3, before mixing with the fibrin solution. The final cell densities produced in hydrogel were 1×10^5^, 1×10^6^, or 1×10^7^ cell/ml. Cell-laden hydrogels were then cultured in MEM for one day before US measurements.

Ultrasound data were acquired as described in section 2.2, and processed as in section 2.1, with the assumptions that the attenuation, speed of sound and density of MEM and DPBS were equal to those of water at 37°C (i.e., *c*_*w*_ = 1524 m/s [21], *ρ*_*w*_ = 993.33 kg/m^3^ [22], and *a* = 1.39×10^−4^ dB/mm/MHz^2^). As for the polystyrene substrate, its speed of sound was set to *c*_*p*_ = 2380 m/s [23], and its density is assumed to be unchanged relative to 22°C (*ρ*_*p*_ = 1050 kg/m^3^) due to the material’s low expansion coefficient. The resulting acoustic impedances for coupling medium and polystyrene were therefore set to *Z*_*w*_ = 1.*5*13 MRayl and *Z*_*p*_ = 2.499 MRayl. The average standard deviations for the thickness and speed of sound across the entire gel ROI area is provided in Supplementary Table 1. The average standard deviation in thickness and speed of sound across the selected 6 mm ROI for an entire gel across all cell conditions was 38 µm and 5 m/s which is <7% of the mean thickness values <1% of the mean speed of sound values (Supplementary Table 1).

### 2.7 Statistical analyses

Results are presented as mean ± standard deviation of the acoustic, physical, or mechanical properties measured over the ROI for each fibrin gel. Measurements were performed in three to six different samples for each cell density and each sample thickness. A one-way ANOVA was used to evaluate the significance of the differences in the means of all measured properties between cell concentrations, with pairwise comparisons using Tukey’s multiple comparisons test (α = 0.001) (GraphPad Prism v8.0, San Diego, USA). Correlations of the speed of sound and acoustic attenuation coefficient with thickness were tested by performing a two-tailed Pearson’s correlation analysis with α = 0.01.

## 3 Results

### 3.1 Acoustic characterization validation

Speed of sound, acoustic impedance, density, and acoustic attenuation coefficient of TPX estimated using the method described in sections 2.1 and 2.2, are compared to those obtained using other measurement methods in Table 1. We found strong agreement between measured and reference values, with a maximum difference of 6.6% (Fig. 2, Table 1). To further validate our technique, we measured the thickness of cell-free fibrin gels in comparison with laser confocal microscopy and found that the thicknesses of cell-free fibrin gels measured by US were highly correlated with those measured by confocal microscopy (r^2^=0.97; p<0.001; Fig. 2C).

**Table 1:** Properties of the TPX film at 22°C (± standard deviation, n=3) compared to published reference values from two studies [19] and [20]. Percent differences for some values are calculated based on the furthest reference value to the measured value from either [19] and/or [20].^§^Acoustic impedance values are presented using two different methods in [20] and thus cannot be directly compared and are not published in [19]. *Acoustic attenuation was calculated via a mean maximum amplitude time domain method and cannot be directly compared with our analytical method. Further discussion of both these discrepancies is presented in the Discussion section.

| Property | Measured value | Reference value | Percent difference | Ref. |
| --- | --- | --- | --- | --- |
| Thickness (mm) | $0.472 \pm 0.004$ | $0.465 \pm 0.001$ | 1.5% | Measured via digital tube micrometer |
| Speed of sound (m/s) | $2180 \pm 14$ @ 22°C | 2093<br>2190 | 4.1%<br>0.5% | [19]<br>[20] |
| Acoustic impedance (MRayl) | $1.70 \pm 0.01$ | $^{\S}1.7, 1.83$ | - | [20] |
| Density (kg/m<sup>3</sup>) | $778 \pm 8$ | 833<br>776 | 6.6 %<br>0.3% | [19]<br>[20] |
| Acoustic attenuation @ 40 MHz (dB/mm) | $8.44 \pm 0.24$ (DX-845 film) | *11.94 (RT-18 film)<br>*9.74 (DX-845 sheet) | - | [19] |

**Figure 2:**
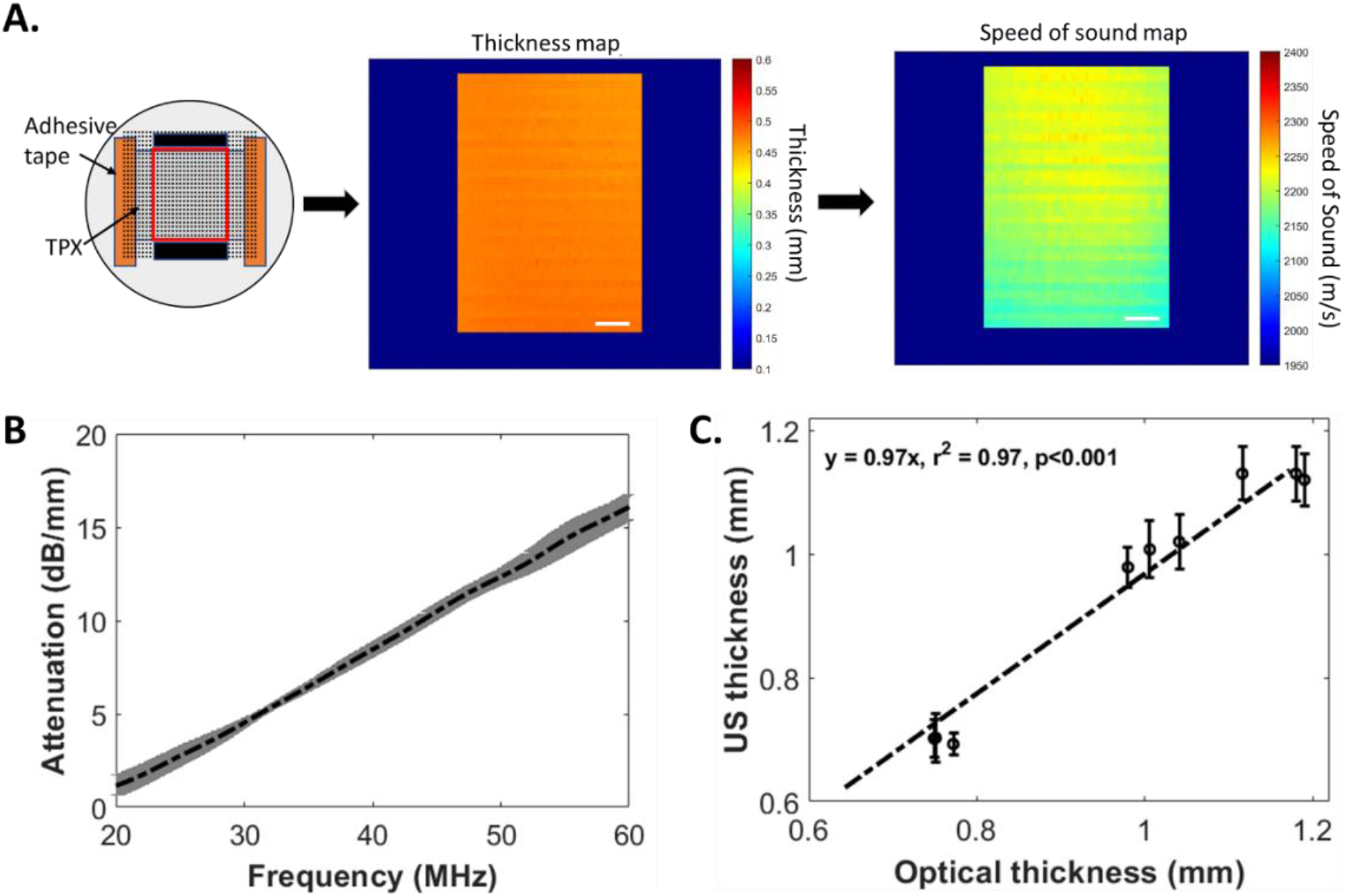
(A-left) TPX film taped down to a polystyrene Petri dish. Dotted lines represent the data acquisition raster scan, with the red line delimiting the ROI used for sample acoustic characterization, and the black rectangles representing the ROIs where the substrate reference signals were collected. (A-middle) estimated local sample thickness. (A-right) local speed of sound. Scale bars = 2 mm. (B) TPX frequency-dependent attenuation. (C) Cell-free fibrin gel thickness: US vs confocal microscopy (error bars: spatial standard deviation of US measurements over the ROI, dashed line: linear fit).

### 3.2 Effects of cell-mediated contraction on physical properties of cell-laden hydrogels

Optical images of cell-laden hydrogels of different thickness and at different cell concentrations are shown on Fig. 3A, showing visual changes in the population of cells in the fibrin hydrogels at each cell condition. Ultrasound measurements showed a spatially uniform shrinkage of the hydrogel associated with cell contraction (Fig. 3B). The standard deviation of the thickness measurements is shown in Supplementary Table 1. This shrinkage increases with cell density, with as much as 60% reduction in the thickness for hydrogels seeded with 1×10^7^ cell/ml relative to their initial thickness (Fig. 3C). Variations in hydrogel thickness, such as those induced by cell contraction, can confound acoustic property measurements if the thickness is not measured accurately in situ. Furthermore, reported acoustic attenuation estimation methods [7-12, 16, 17], which did not compensate for local losses associated with reflection and transmissions at interfaces, introduce a thickness-dependent bias *b*, in the estimated attenuation coefficient, which is expressed as:

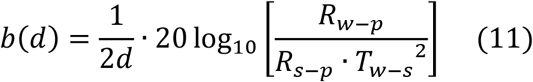

**Figure 3:**
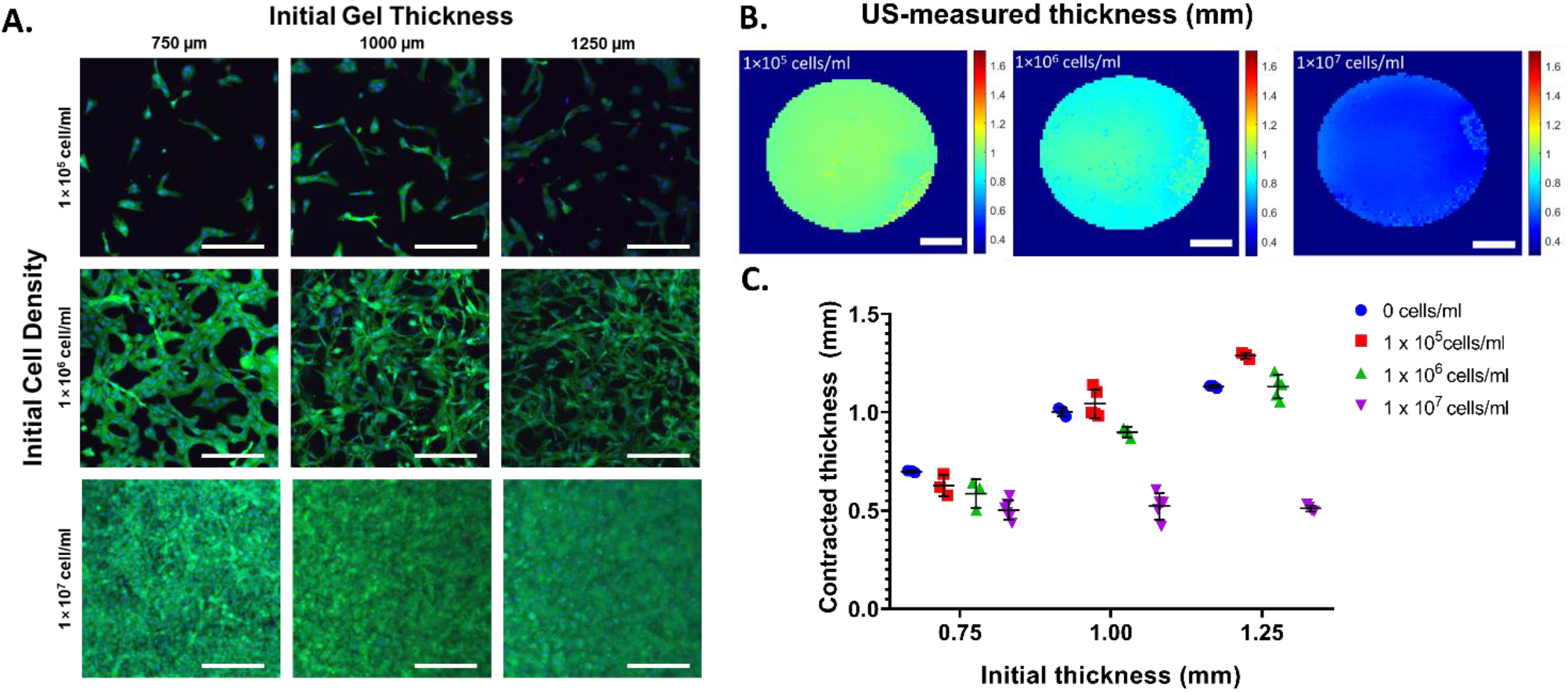
(A) Optical images of cell-laden hydrogels of three different initial thicknesses (columns) and at three different cell densities (rows). Live cells, green; dead cells, red; nucleus, blue. Scale bars = 200 µm. (B) US-measured thicknesses of hydrogels of 1.00 mm initial thickness, with cell densities of, from left to right, 1×10^5^, 1×10^6^ and 1×10^7^ cell/ml. Scale bars = 1 mm. (C) Effect of cell contraction on hydrogel thickness measured via US compared to the initial hydrogel thickness.

This bias can become significant for small sample thicknesses and is presented for all measured samples in Supplementary Table 2.

To evaluate whether the method could overcome the aforementioned limitations, we investigated the thickness dependence of the acoustic properties of cell-laden hydrogels. Results summarized in Fig. 4 demonstrate that speed of sound, acoustic attenuation, density, and elastic modulus were not significantly correlated with gel thickness over a wide range of cell densities and gel thicknesses (Fig. 4; p>0.11, p>0.23, p>0.22, p>0.05 across cell densities via Pearson’s, respectively). Acoustic impedance was only statistically correlated with thickness in the 1×10^6^ cell/ml condition (p<0.01, r = -0.85), but with weak dependency as values varied by <0.2% across seven orders of magnitude of cell density (corresponding to minor absolute differences of ∼0.002 MRayl).

**Figure 4:**
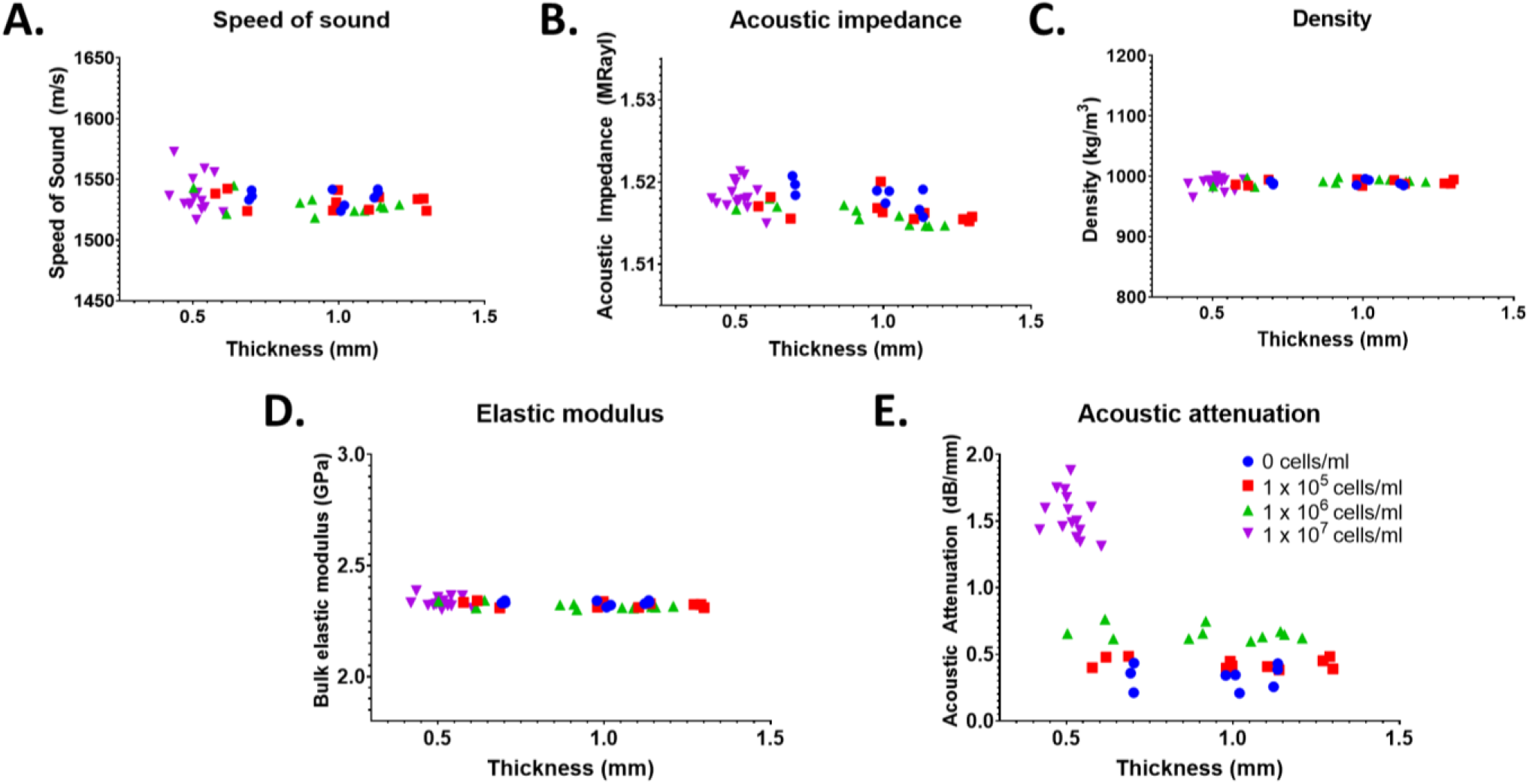
Thickness dependence of (A) speed of sound, (B) acoustic impedance, (C) acoustic attenuation, (D) density, and (E) elastic modulus. None of the properties were significantly correlated with hydrogel thickness (p>0.01; Pearson’s), except for a weak correlation for acoustic impedance at 1×10^6^ cell/ml (p<0.01; r = -0.85).

### 3.3 Acoustic properties of cell-laden hydrogels

Since thickness was determined to be a non-confounding factor in our method (Fig. 4), cell density in these cell-laden hydrogels can be considered the primary factor driving affecting the acoustic attenuation coefficient. The acoustic properties (e.g., speed of sound, acoustic impedance, acoustic attenuation coefficient), density, and elastic modulus of cell-laden gels (n=46) are reported as a function of initial cell density in Figure 5 (summarized in Supplementary Table 1). There were no statistically significant difference in the speed of sound across cell densities, with <0.6% difference between the mean values (p=0.25; Fig. 5A). Similarly, acoustic impedance measurements varied by <0.2% across cell densities, although a statistical difference was detected between the two highest cell densities (Fig. 5B). Since speed of sound and impedance are mostly invariant with cell density in the fibrin gels, the derived density and elastic modulus varied minimally across cell densities. The mean densities of the fibrin hydrogels (988 – 991 kg/m) were approximately those of water [21], with no significant differences across cell densities (Fig. 5C; p>0.59). Similarly, elastic modulus measurements varied by less than <0.7% across cell densities (corresponding to absolute differences of <0.02 GPa), with no significant differences across cell densities (Fig. 5D; p>0.08). In contrast, the acoustic attenuation coefficient measured at 40 MHz increased significantly with increasing cell density (Fig. 5E; p<0.001), except between cell-free and 1×10^5^ cell/ml. This increase is observed across the bandwidth of the transducer (20 – 60 MHz) as shown in Figure 5F.

**Figure 5:**
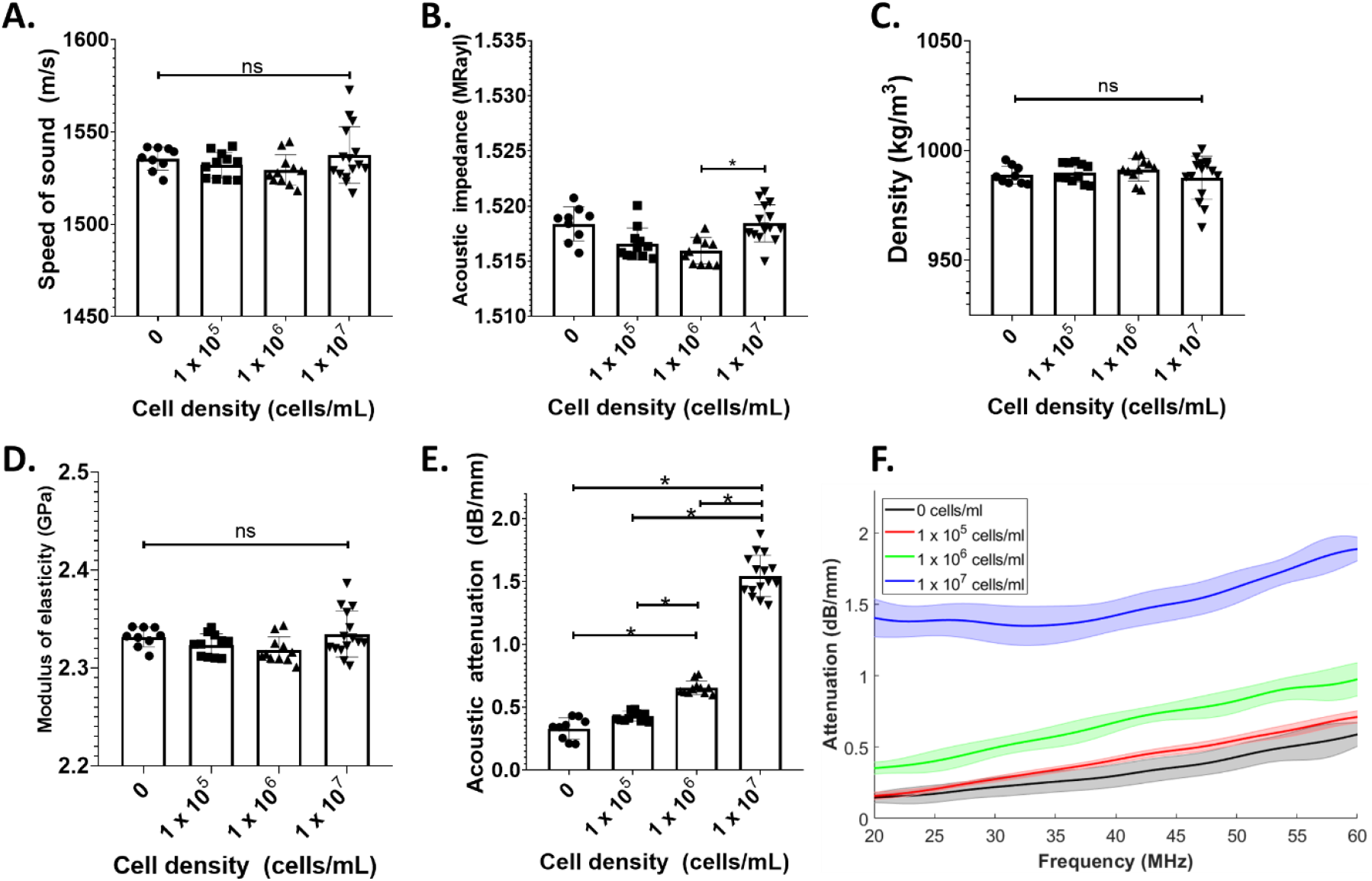
Measurements of the acoustic properties of cell-laden hydrogels at four different cell densities and three thicknesses, including (A) speed of sound and (B) acoustic impedance via our ultrasound method, (C) density and (D) modulus of elasticity derived from the values in (A-B), and (E) acoustic attenuation at 40 MHz. (F) Representative frequency-dependent attenuation plot for each cell density in a 1.00 mm gel within the bandwidth of the transducer (20-60 MHz). Standard deviations at each frequency are represented as shaded areas around the mean attenuation. A minimum of n=3 independent gels were used per thickness for each initial cell density. Non-significant (ns) statistical differences between groups by a one-way ANOVA are shown and significant differences from a post-hoc Tukey’s multiple comparison test are shown with *p < 0.001.

## 4 Discussion

US has excellent potential for non-invasive, real-time monitoring of the acoustic, physical, and mechanical properties of biomaterials and engineered tissues. Here, we developed an analytical approach that addresses the limitations of previous approaches and validated it by measuring the acoustic properties of TPX. We then applied this technique in cell-laden fibrin hydrogels. Importantly, our approach accurately measured hydrogel thickness, despite significant cell-mediated contraction, which enabled estimations of acoustic properties that were effectively independent of thickness and reflected intrinsic hydrogel properties.

Since sample thickness plays a critical role in evaluating the acoustic properties, the acoustic measurement of the thickness of TPX and fibrin hydrogel samples was compared to a digital tube micrometer and a laser confocal microscopy measurement, respectively, and good agreement was found. To validate our acoustic measurement technique, we compared the physical and acoustic properties of TPX at 40 MHz obtained using our analytical approach to those obtained with the methods used by Madsen et al [19] and Bloomfield et al [20]. The speed of sound determined via our method was within range of the speed determined in [19] and [20]. Bloomfield et al. [20] present two acoustic impedance values for TPX for two different measurement methods. Using reflection coefficient measurements, Bloomfield et al. also calculate an acoustic impedance of TPX of 1.7 MRayl which is consistent with our measurements (Table 1). However, using a measured speed of sound and a manufacturer-provided density, Bloomfield et al. determine the acoustic impedance of TPX to be 1.83 MRayl. The latter method is less accurate as it relies on a non-experimentally derived value but was incorrectly used for density calculations. Instead of using the manufacturer-provided density, if Bloomfield et al. had used the pulse-echo-measured impedance (1.7 MRayl) and their measured speed of sound (2190 m/s) to estimate the density of TPX, their obtained value would have been 776 kg/m^3^ which is within <1% of our study’s measured density (Table 1). With regards to attenuation, we found a difference between our measurements and those from [19], and [20], which is most likely due to the difference in measurement approaches used in each study. One should note that in materials for which attenuation is highly dependent on frequency, such as TPX, broad-band time domain amplitude measurement methods ([19, 20]) are suboptimal due to downshift of the central frequency of the ultrasound pressure pulse signal propagating through the sample. Moreover, from Eq.11, there is a bias of 2.58 dB/mm when local losses associated with reflection and transmissions at the interfaces are not accounted for. Through addition of this bias to our measurement of the acoustic attenuation of TPX at 40 MHz of 8.44 dB/mm, we are within range of [19]. Nevertheless, the acoustic attenuation measured by our largely frequency-based approach should be more accurate than [19] across different sample types because it accounts for changes in the peak frequency of the media rather than the ratio of the amplitude of the time-domain signals.

Of the acoustic properties measured, the acoustic attenuation coefficient had the most notable change with increased cell density. Our acoustic attenuation coefficient values are within range of other published studies [8, 9, 16, 17], which have not accounted for all physical interfacial phenomena. The average bias for cell-laden hydrogels is 0.03 dB/mm (Supplementary Table 2) because the acoustic impedance of the fibrin hydrogels is similar to that of the coupling liquid. However, for: 1) thicker engineered tissues or implantable biomaterials, this bias would be prominent; and 2) for stiffer engineered tissues (e.g., cartilage, bone), this acoustic impedance mismatch would be higher and cause the bias to have more significant effects. As the fibrin hydrogels were >99% water, the measured speed of sound, derived density, and derived bulk modulus of the gels were approximately that of water at 37°C (1524 m/s, 993.33 kg/m^3^, 2.32 Gpa from Eq. 8, respectively). We did not expect the addition of cells within the fibrin hydrogels to affect the speed of sound, acoustic impedance, density, and elastic modulus (assuming no shear propagation) of the cell-laden fibrin hydrogels since cells are largely (≥70% [25]) composed of water with a high concentration of salt and have properties that approach those of 37°C water [26]. In contrast, acoustic impedance increased with cell density ostensibly due to absorption and scattering by the cells’ lipidic membranes [27] and nuclei [14], respectively.

Our study has limitations: (1) Our method calculates acoustic impedance using the reflection coefficient at the water-sample interface as a ratio between the pulse-echo signal from the water-substrate interface and the water-sample interface. As such, it is sensitive to local variations in the interface topography, e.g., due to focal cell topography, contraction, or swelling. (2) Non-uniform distribution of cells through the thickness of the fibrin hydrogels could lead to underestimation of the attenuation values, as Eq.1 is normalized to the thickness of the entire gel (*d*) rather than to the thickness of where the dominant population of cells are located during the actual measurement. (3) Lastly, although our technique is versatile and can be used with any substrate material of know acoustic properties, it requires good adherence of the sample to the substrate to avoid air gaps between the sample and substrate. However, regardless of the substrate and/or sample material, the frequency-based approach presented in this study can be used to estimate the acoustic properties of cell-laden hydrogels across multiple thicknesses and cell densities. In future work, the approach could be used to monitor dynamics changes with culture time.

## 5 Conclusion

Here, we present a non-invasive and non-destructive high-frequency quantitative US method to estimate the acoustic, physical, and mechanical properties of biomaterials and engineered tissues with high resolution (∼100 µm). This method accounts for all physical interfacial phenomena to estimate the intrinsic physical and acoustic properties of cell-laden hydrogels, including thickness, speed of sound, acoustic impedance, acoustic attenuation coefficient, density, and elastic modulus, without the confounding variable of hydrogel thickness. We showed that increased cell density in the hydrogel led to an increase in the acoustic attenuation coefficient, demonstrating the ability of high-frequency US to detect changes in the cellularity of cell-laden biomaterials. Importantly, the method corrects for changes in hydrogel thickness due to cell-mediated contraction. This robust technique enables non-invasive, non-destructive estimation of the acoustic, physical, and mechanical properties of cell-laden biomaterials at spatial resolutions that are relevant for tissue engineering applications.

## 6 Acknowledgements

This work was supported by a Collaborative Health Research Program grant from the Natural Science and Engineering Research Council of Canada (NSERC; CHRPJ 5083) and Canadian Institutes of Health Research (CPG-151946). J.A.S was supported by an NSERC Canada Graduate Scholarship – Master’s, a CRAFT Doctoral Fellowship, an NSERC CREATE Doctoral Fellowship, and an NSERC Vanier Canada Graduate Scholarship. E.M.S was supported by an NSERC postdoctoral scholarship and a Medicine by Design Postdoctoral Fellowship. B.M. was supported by an NSERC Canada Graduate Scholarship – Doctoral.

## 7 Data availability

The raw data and codes to process the raw data required to reproduce these findings of this study are available to download from <u>here</u>.

## References

[1] Dalecki, D., & Hocking, D. C. (2016). Advancing Ultrasound Technologies for Tissue Engineering. In Handbook of Ultrasonics and Sonochemistry (pp. 1101–1126). Springer Singapore. https://doi.org/10.1007/978-981-287-278-4_28

[2] Kim, K., & Wagner, W. R. (2016). Non-invasive and Non-destructive Characterization of Tissue Engineered Constructs Using Ultrasound Imaging Technologies: A Review. Annals of Biomedical Engineering, 44(3), 621–635. https://doi.org/10.1007/s10439-015-1495-0

[3] Dalecki, D., Mercado, K. P., & Hocking, D. C. (2016). Quantitative Ultrasound for Nondestructive Characterization of Engineered Tissues and Biomaterials. Annals of Biomedical Engineering, 44(3), 636–648. https://doi.org/10.1007/s10439-015-1515-0

[4] Deng, C. X., Hong, X., & Stegemann, J. P. (2016). Ultrasound Imaging Techniques for Spatiotemporal Characterization of Composition, Microstructure, and Mechanical Properties in Tissue Engineering. Tissue Engineering Part B: Reviews, 22(4), 311–321. https://doi.org/10.1089/ten.teb.2015.0453

[5] Dalecki, D., & Hocking, D. C. (2015). Ultrasound Technologies for Biomaterials Fabrication and Imaging. Annals of Biomedical Engineering, 43(3), 747–761. https://doi.org/10.1007/s10439-014-1158-6

[6] Aliabouzar, M., Zhang, G. L., & Sarkar, K. (2018). Acoustic and mechanical characterization of 3D-printed scaffolds for tissue engineering applications. Biomedical Materials, 13(5), 055013. https://doi.org/10.1088/1748-605X/aad417

[7] Cafarelli, A., Verbeni, A., Poliziani, A., Dario, P., Menciassi, A., & Ricotti, L. (2017). Tuning acoustic and mechanical properties of materials for ultrasound phantoms and smart substrates for cell cultures. Acta Biomaterialia, 49, 368–378. https://doi.org/10.1016/j.actbio.2016.11.049

[8] Mercado, K. P., Helguera, M., Hocking, D. C., & Dalecki, D. (2015). Noninvasive Quantitative Imaging of Collagen Microstructure in Three-Dimensional Hydrogels Using High-Frequency Ultrasound. Tissue Engineering Part C: Methods, 21(7), 671–682. https://doi.org/10.1089/ten.tec.2014.0527

[9] Ruland, A., Gilmore, K. J., Daikuara, L. Y., Fay, C. D., Yue, Z., & Wallace, G. G. (2019). Quantitative ultrasound imaging of cell-laden hydrogels and printed constructs. Acta Biomaterialia, 91, 173–185. https://doi.org/10.1016/j.actbio.2019.04.055

[10] Gudur, M. S. R., Rao, R. R., Peterson, A. W., Caldwell, D. J., Stegemann, J. P., & Deng, C. X. (2014). Noninvasive Quantification of In Vitro Osteoblastic Differentiation in 3D Engineered Tissue Constructs Using Spectral Ultrasound Imaging. PLoS ONE, 9(1), e85749. https://doi.org/10.1371/journal.pone.0085749

[11] Mercado, K. P., Helguera, M., Hocking, D. C., & Dalecki, D. (2014). Estimating Cell Concentration in Three-Dimensional Engineered Tissues Using High Frequency Quantitative Ultrasound. Annals of Biomedical Engineering, 42(6), 1292–1304. https://doi.org/10.1007/s10439-014-0994-8

[12] D’Astous, F. T., & Foster, F. S. (1986). Frequency dependence of ultrasound attenuation and backscatter in breast tissue. Ultrasound in Medicine & Biology, 12(10), 795–808. https://doi.org/10.1016/0301-5629(86)90077-3

[13] Strohm, E. M., Czarnota, G. J., & Kolios, M. C. (2010). Quantitative measurements of apoptotic cell properties using acoustic microscopy. IEEE Transactions on Ultrasonics, Ferroelectrics and Frequency Control, 57(10), 2293–2304. https://doi.org/10.1109/TUFFC.2010.1690

[14] Taggart, L. R., Baddour, R. E., Giles, A., Czarnota, G. J., & Kolios, M. C. (2007). Ultrasonic Characterization of Whole Cells and Isolated Nuclei. Ultrasound in Medicine & Biology, 33(3), 389–401. https://doi.org/10.1016/j.ultrasmedbio.2006.07.037

[15] Vernerey, F. J., Lalitha Sridhar, S., Muralidharan, A., & Bryant, S. J. (2021). Mechanics of 3D Cell–Hydrogel Interactions: Experiments, Models, and Mechanisms. Chemical Reviews, 121(18), 11085–11148. https://doi.org/10.1021/acs.chemrev.1c00046

[16] Ruland, A., Hill, J. M., & Wallace, G. G. (2021). Reference Phantom Method for Ultrasonic Imaging of Thin Dynamic Constructs. Ultrasound in Medicine & Biology, 47(8), 2388–2403. https://doi.org/10.1016/j.ultrasmedbio.2021.04.014

[17] Ruland, A., Onofrillo, C., Duchi, S., Di Bella, C., & Wallace, G. G. (2022). Standardised quantitative ultrasound imaging approach for the contact-less three-dimensional analysis of neocartilage formation in hydrogel-based bioscaffolds. Acta Biomaterialia, 147, 129–146. https://doi.org/10.1016/j.actbio.2022.05.037

[18] Briggs, G. A. D., Wang, J., & Gundle, R. (1993). Quantitative acoustic microscopy of individual living human cells. Journal of Microscopy, 172(1), 3–12. https://doi.org/10.1111/j.1365-2818.1993.tb03387.x

[19] Madsen, E. L., Deaner, M. E., & Mehi, J. (2011). Properties of Phantom Tissuelike Polymethylpentene in the Frequency Range 20–70 MHZ. Ultrasound in Medicine & Biology, 37(8), 1327–1339. https://doi.org/10.1016/j.ultrasmedbio.2011.05.023

[20] Bloomfield, P. E., Wei-Jung Lo, & Lewin, P. A. (2000). Experimental study of the acoustical properties of polymers utilized to construct PVDF ultrasonic transducers and the acousto-electric properties of PVDF and P(VDF/TrFE) films. IEEE Transactions on Ultrasonics, Ferroelectrics and Frequency Control, 47(6), 1397–1405. https://doi.org/10.1109/58.883528

[21] Del Grosso, V. A., & Mader, C. W. (1972). Speed of Sound in Pure Water. The Journal of the Acoustical Society of America, 52(5B), 1442–1446. https://doi.org/10.1121/1.1913258

[22] Haynes, W. M. (2014). CRC handbook of chemistry and physics: A ready-reference book of chemical and physical data.

[23] Kono, R. (1960). The Dynamic Bulk Viscosity of Polystyrene and Polymethyl Methacrylate. Journal of the Physical Society of Japan, 15(4), 718–725. https://doi.org/10.1143/JPSJ.15.718

[24] Treeby, B. E., Zhang, E. Z., Thomas, A. S., & Cox, B. T. (2011). Measurement of the Ultrasound Attenuation and Dispersion in Whole Human Blood and its Components From 0–70 MHz. Ultrasound in Medicine & Biology, 37(2), 289–300. https://doi.org/10.1016/j.ultrasmedbio.2010.10.020

[25] Cooper GM. The Cell: A Molecular Approach. 2nd edition. Sunderland (MA): Sinauer Associates; 2000. The Molecular Composition of Cells.

[26] Wirtzfeld, L. A., Berndl, E. S. L., Czarnota, G. J., & Kolios, M. C. (2017). Monitoring Quantitative Ultrasound Parameter Changes in a Cell Pellet Model of Cell Starvation. Biophysical Journal, 112(12), 2634–2640. https://doi.org/10.1016/j.bpj.2017.05.017

[27] Sundaram, J., Mellein, B. R., & Mitragotri, S. (2003). An Experimental and Theoretical Analysis of Ultrasound-Induced Permeabilization of Cell Membranes. Biophysical Journal, 84(5), 3087–3101. https://doi.org/10.1016/S0006-3495(03)70034-4

