## Supplementary Information for "High-frequency quantitative ultrasound to assess the acoustic properties of engineered tissues in vitro"

### Supplementary Information for:
### High-frequency quantitative ultrasound to assess the acoustic properties of engineered tissues in vitro

**Authors:** Joseph A. Sebastian[1,2], Eric M. Strohm[3,4,5], Emmanuel Chérin[6], Bahram Mirani[1,2,7], Christine Démoré[6,8], Michael C. Kolios[3,4,5], Craig A. Simmons[1,2,7]

**Affiliations:**

1. Institute of Biomedical Engineering, University of Toronto, Toronto, Canada
2. Translational Biology and Engineering Program, Ted Rogers Center for Heart Research, Toronto, Canada
3. Department of Physics, Toronto Metropolitan University, Toronto, Canada
4. Institute of Biomedical Engineering, Science and Technology (iBEST), a partnership between Toronto Metropolitan University and St. Michael's Hospital, Toronto, Canada
5. Keenan Research Centre for Biomedical Science, Li Ka Shing Knowledge Institute, St. Michael's Hospital, Toronto, Canada
6. Sunnybrook Research Institute, Toronto, Canada
7. Department of Mechanical and Industrial Engineering, University of Toronto, Toronto, Canada
8. Department of Medical Biophysics, University of Toronto, Toronto, Canada

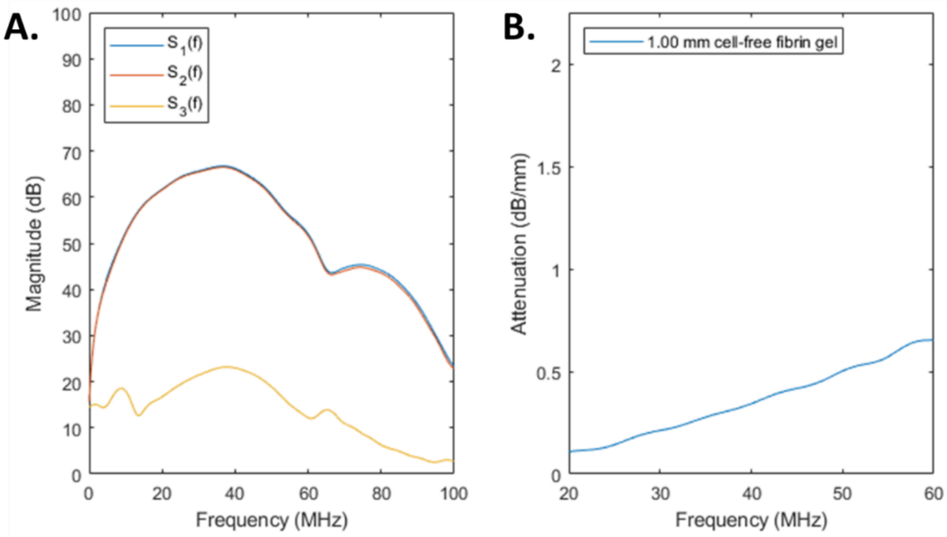


**Supplementary Figure 1:** (A) Representative frequency-domain signals for spectral echoes at the liquid-substrate interface ($S_1(f)$), sample-substrate interface ($S_2(f)$), liquid-sample interface ($S_3(f)$) for a 1.00 mm cell-free fibrin gel and (B) Representative frequency-dependent attenuation curve for the spectral echoes in (A) determined using Equation 1.

**Supplementary Table 1**: Average standard deviation of thickness and speed of sound values across entire gel area for three thicknesses per cell condition.

| Cell density (cells/mL) | Average standard deviation of US-measured thickness (µm) | Average standard deviation of speed of sound (m/s) |
|---|---|---|
| 0 | 38 | 5.9 |
| $1 \times 10^5$ | 40 | 5.4 |
| $1 \times 10^6$ | 41 | 3.4 |
| $1 \times 10^7$ | 35 | 7.4 |

**Supplementary Table 2**: Means and standard deviation of all measured samples grouped by cell density.

| Cell density (cells/mL) | Speed of sound (m/s) | Acoustic impedance (MRayl) | Acoustic attenuation (dB/mm) | Bias of acoustic attenuation (dB/mm) | Density (kg/m$^3$) | Elastic modulus (GPa) |
|---|---|---|---|---|---|---|
| 0 | 1536 ± 6 | 1.518 ± 0.002 | 0.330 ± 0.086 | 0.030 ± 0.014 | 988.8 ± 4.1 | 2.332 ± 0.010 |
| $1 \times 10^5$ | 1532 ± 7 | 1.517 ± 0.001 | 0.431 ± 0.039 | 0.017 ± 0.011 | 989.9 ± 4.3 | 2.324 ± 0.011 |
| $1 \times 10^6$ | 1529 ± 8 | 1.516 ± 0.001 | 0.657 ± 0.053 | 0.015 ± 0.011 | 991.3 ± 5.2 | 2.318 ± 0.013 |
| $1 \times 10^7$ | 1538 ± 15 | 1.518 ± 0.002 | 1.546 ± 0.164 | 0.050 ± 0.018 | 987.7 ± 9.8 | 2.335 ± 0.024 |
